# Distinctive types of postzygotic single-nucleotide mosaicisms in healthy individuals revealed by genome-wide profiling of multiple organs

**DOI:** 10.1101/309005

**Authors:** August Yue Huang, Xiaoxu Yang, Sheng Wang, Xianing Zheng, Qixi Wu, Adam Yongxin Ye, Liping Wei

## Abstract

Postzygotic single-nucleotide mosaicisms (pSNMs) have been extensively studied in tumors and are known to play critical roles in tumorigenesis. However, the patterns and origin of pSNMs in normal organs of healthy humans remain largely unknown. Using whole-genome sequencing and ultra-deep amplicon re-sequencing, we identified and validated 164 pSNMs from 27 postmortem organ samples obtained from five healthy donors. The mutant allele fractions ranged from 1.0% to 29.7%. Inter- and intra-organ comparison revealed two distinctive types of pSNMs, with about half originating during early embryogenesis (embryonic pSNMs) and the remaining more likely to result from clonal expansion events that had occurred more recently (clonal expansion pSNMs). Compared to clonal expansion pSNMs, embryonic pSNMs had higher proportion of C>T mutations with elevated mutation rate at CpG sites. We observed differences in replication timing between these two types of pSNMs, with embryonic and clonal expansion pSNMs enriched in early- and late-replicating regions, respectively. An increased number of embryonic pSNMs were located in open chromatin states and topologically associating domains that transcribed embryonically. Our findings provide new insights into the origin and spatial distribution of postzygotic mosaicism during normal human development.

**Author Summary:** Genomic mosaicism led by postzygotic mutation is the major cause of cancers and many non-cancer developmental disorders. Theoretically, postzygotic mutations should be accumulated during the developmental process of healthy individuals, but the genome-wide characterization of postzygotic mosaicisms across many organ types of the same individual remained limited. In this study, we identified and validated two types of postzygotic mosaicism from the whole-genomes of 27 organs obtained from five healthy donors. We further found that the postzygotic mosaicisms arising during early embryogenesis and later clonal expansion events show distinct genomic patterns in mutation spectrum, replication timing, and chromatin status.

## Introduction

Postzygotic mutations refer to DNA changes arising after the formation of the zygote that lead to genomic mosaicisms in a single individual [1, 2]. Unlike *de novo* or inherited germline mutations, postzygotic mutations only affect a fraction of cells in multicellular organisms, and individuals carrying a functional mosaic mutation typically exhibit a milder phenotype [3–5]. The roles of postzygotic single-nucleotide mosaicisms (pSNMs) have been demonstrated in numerous cancers [6, 7] and various types of developmental disorders, including malformations [8, 9] and autism [10, 11]. We and another research group have reported the first genome-wide identification and characterization of pSNMs from the peripheral blood samples of healthy individuals [12, 13]. More recently, the accumulation of postzygotic mutations during aging process has been reported in blood or brain samples [14–17]. Yadav *et al.* studied pSNMs in apparently benign tissue samples obtained from cancer patients [18], but the contribution of pre-cancerous mutations could not be completely ruled out and the study was restricted to exonic regions. As such, the occurrence and genomic pattern of pSNMs in normal tissues of healthy individuals remains under-investigated.

It has been reported that cancer genomes have distinct mutational signatures resulting predominantly from exposure to mutagenic agents and dysfunction of the DNA repair machinery [19]. Additional genomic factors, such as replication timing and chromatin status, could also impact the distribution of pSNMs in cancer genomes [20–22]. Whether and how these genomic factors might contribute to the genomic distribution of pSNMs in organs of healthy individuals remains largely unexplored [23]. Tumorigenesis has been considered as an evolutionary process in which tumor cells with increased fitness will proliferate faster than normal cells and lead to the clonal expansion of tumor cell population in a specific organ [24, 25]. Although such events of clonal expansion have been previously reported in apparently normal skin and blood samples [17, 26], it remains unclear whether clonal expansion plays a role in other non-cancer tissue types. Understanding the origin and spatial distribution of pSNMs in normal tissues of healthy individuals could provide an important baseline for interpreting their contributions to disease states [27].

The next-generation sequencing technologies (NGS) have greatly advanced the study of pSNMs [28]. Sequencing the genomes of single cells after whole-genome amplification or *in vivo* clonal proliferation have been applied to the study of pSNM profiles of normal human cells, including germ cells [29], adult stem cells [30], and neurons [31]. Typically, tens or hundreds of cells from each sample need to be sequenced to identify and quantify pSNMs, which tends to increase the cost [32]. The inaccurate process of whole-genome amplification in single-cell sequencing makes it difficult to distinguish real pSNMs from technical artifacts, and the challenge of rigorously validating the pSNMs in a cell that have been already amplified aggravates the uncertainties [33, 34]. Bulk sequencing is potentially a reliable and cost-effective alternative that, importantly, allows for rigorous validations of pSNMs [23]. Utilizing bulk whole-genome sequencing (WGS) and ultra-deep amplicon re-sequencing, this current study identified and validated pSNMs from 27 different organ samples obtained from five healthy donors and investigated the origin and spatial distribution of pSNMs in the developmental process of these healthy individuals.

## Results

### Genome-wide Identification and Profiling of Postzygotic Single-Nucleotide Mosaicisms in Healthy Human Organ Samples

Postmortem organ samples from five healthy Asian donors (age 20-45 yr) were obtained from BioServe, including a total of 27 organ samples from brain, liver, colon, skin, artery, breast, ovary, and prostate (Table 1). The donors died from motor vehicle accidents and were not known to be affected by any types of cancer or other overgrowth disorders. The samples were sequenced using an Illumina HiSeq X Ten sequencing platform with an average depth of 114-150X (Table 1).

**Table 1.**
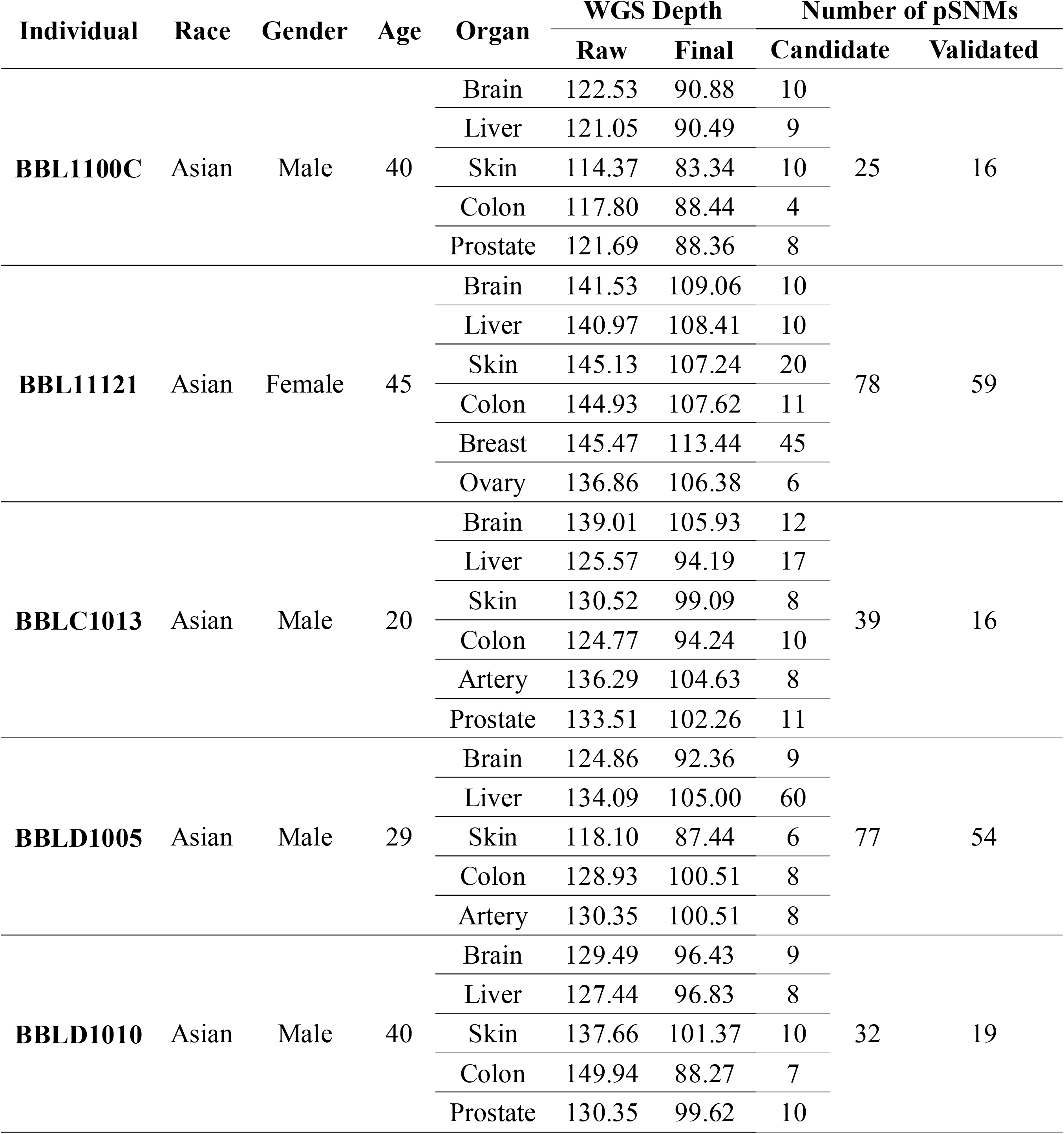
Donor and sequencing information regarding the 27 post-mortem organ samples profiled in this study.

It is expected that many postzygotic mutations occurring at an early stage of embryogenesis may be shared between two or more organs from one individual [13, 27]. Thus, conventional mutation callers, which require matched negative control samples for comparison, would likely miss these mutations. We had previously developed MosaicHunter [35], a bioinformatics pipeline that can detect pSNMs without the need of control sample obtained from the same individual. MosaicHunter incorporated a Bayesian genotyper to distinguish pSNMs from germline variants and base-calling errors and a series of stringent filters to remove systematic errors. Using the Bayesian genotyper, we calculated the posterior probability of mosaic genotype versus three germline genotypes across all the genomic sites with at least 5% mutant allele fraction (the fraction of reads supporting the mutant allele) and 3 or more reads supporting the mutant allele. As a result, we identified a total of 251 candidate pSNMs in the 27 samples from the five donors; among them 41 pSNMs were found in more than one sample from the same donor (Table 1).

Next, we validated the pSNMs and quantified their minor allele fractions in all of the organ samples (Methods). We used an amplicon-based ultra-deep resequencing method, PASM (<u>P</u>GM <u>A</u>mplicon <u>S</u>equencing of <u>M</u>osaicism), which we had previously developed and benchmarked [5]. Of the 251 candidate pSNM sites, 27 were excluded due to failure to design amplicon primers or to get enough sequencing depth in the negative controls. For the remaining 224 sites, the average sequencing depth of the amplicons was greater than 4000X per sample (**S1 Fig**). The peripheral blood samples of two unrelated healthy Asians (ACC1 and ACC4) served as negative controls. A pSNM was considered validated only if the mutant allele was detected in an organ sample in a mosaic state but undetectable in both the negative controls. Three sites with abnormal copy numbers estimated from the WGS data were further excluded (**S1 Table** and Methods). In summary, we successfully validated 164 pSNMs in these five donors, with an overall validation rate of 73.2% (Table 1). The full list of the 164 validated pSNMs was described in **S2 Table**, which was used in the following analyses.

The validated pSNMs were located in 21 autosomes and the X chromosome (Fig 1A). We calculated the genomic distance between nearby pSNMs and found no significant difference between the observed and expected distances if pSNMs were uniformly distributed along the human genome (Kolmogorov-Smirnov test, *P*-value > 0.05), indicating that there was no observable clustering of postzygotic mutations in healthy individuals. The minor allele fractions estimated by PASM ranged from 1.0% to 29.7%, significantly correlating with the fractions estimated by WGS (Fig 1B; Pearson’s *r* = 0.89 and *P*-value < 2.2×10^-16^). The allele fraction of each pSNM varied across the different organs from the same donor (**S2-S6 Figs**).

**Fig 1.**
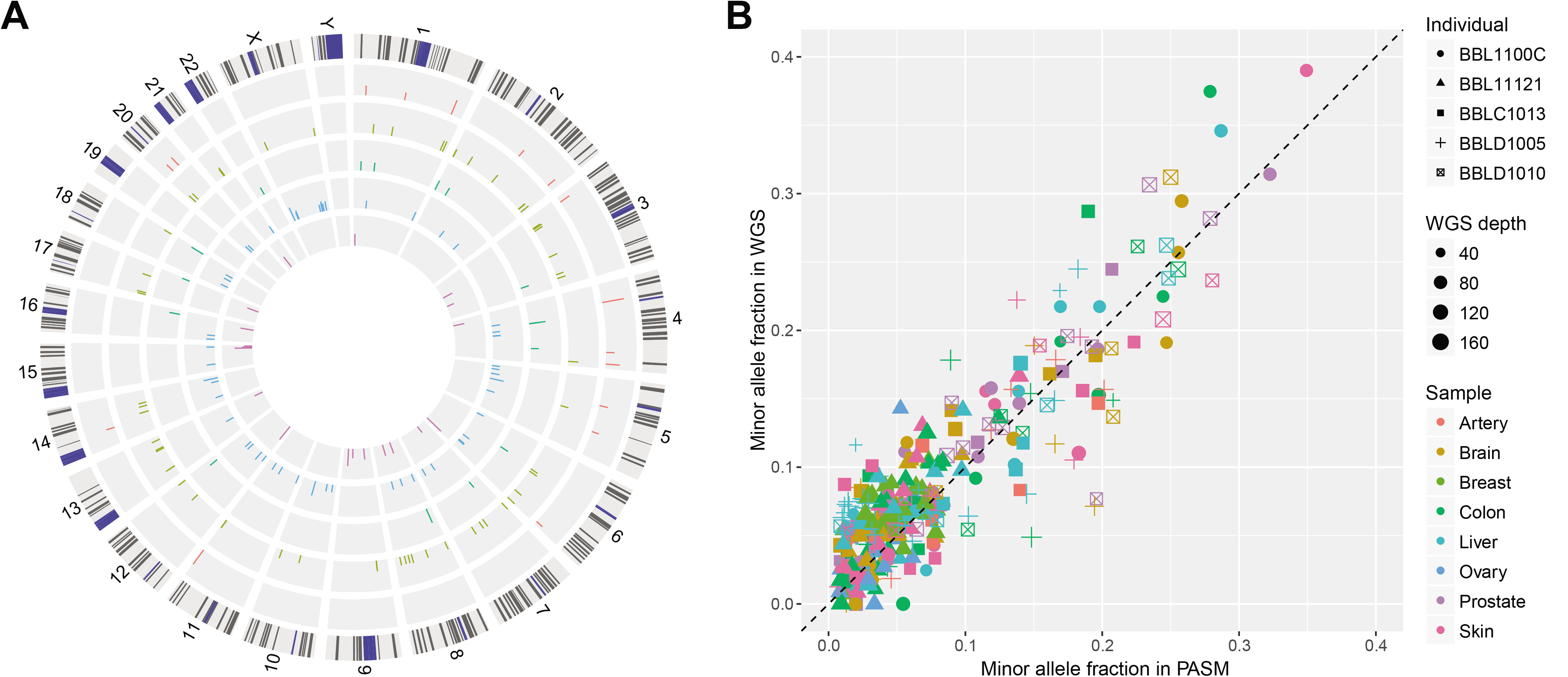
Identification and validation of pSNMs in 27 organ samples obtained from five individuals. (A) Genomic landscape of the validated pSNMs. Circos plots from outer to inner represent individuals BBL1100C, BBL11121, BBLC1013, BBLD1005, and BBLD1010, respectively. The Y axis denotes the average minor allele fraction across pSNM-carrying organs of each donor. (B) Correlation of the allele fractions estimated by whole-genome sequencing and targeted ultra-deep resequencing (PASM) of the validated sites. The shape, color, and size of the dots represent the donor and organ type carrying the pSNMs and the site-specific depth of whole-genome sequencing.

### Two Distinct Types of Postzygotic Single-Nucleotide Mosaicisms Revealed by Inter- and Intra-organ Comparisons

Based on the presence or absence of the validated pSNMs in the organ samples from an individual donor, we grouped the pSNMs into two categories: 60 were present in two or more organ samples of the same donor (27 of which were globally present in all the sequenced organs of the donor) and 104 were uniquely present in a single organ. Given the low postzygotic mutation rate in healthy individuals [36], it was unlikely that multiple postzygotic mutation events involving the same nucleotide alteration occurred independently within one individual. It was more likely that the pSNMs shared by more than one organ resulted from mutation events that had occurred at early developmental stages, and the mutant alleles were passed on to cell lineages of more than one organ type. Comparison of the minor allele fractions in the two categories of pSNMs supported this hypothesis. As shown in Fig 2A, the allele fraction of pSNMs shared by more than one organ was significantly higher than that of pSNMs unique to only one organ (Wilcoxon rank-sum test, *P*-value = 1.2×10^-3^). In particular, 40% (24 out of 60) of the pSNMs shared by more than one organ had allele fractions greater than 1/16, suggesting that they might have originated during the first few cell divisions of embryogenesis [37]. We refer to these pSNMs shared by more than one organ as “embryonic pSNMs” in the following analyses. On average, we identified 4.6~14.5 embryonic pSNMs from each organ of the five individuals, and the occurrence rate was similar across different organs (Fig 2B). We further compared the allele fractions across multiple organs of the same individual, and found that more than 95% of the embryonic pSNMs showed <5% standard deviation of allele fraction (**S2 Table**), indicating no dramatic allele fraction change for embryonic pSNMs.

**Fig 2.**
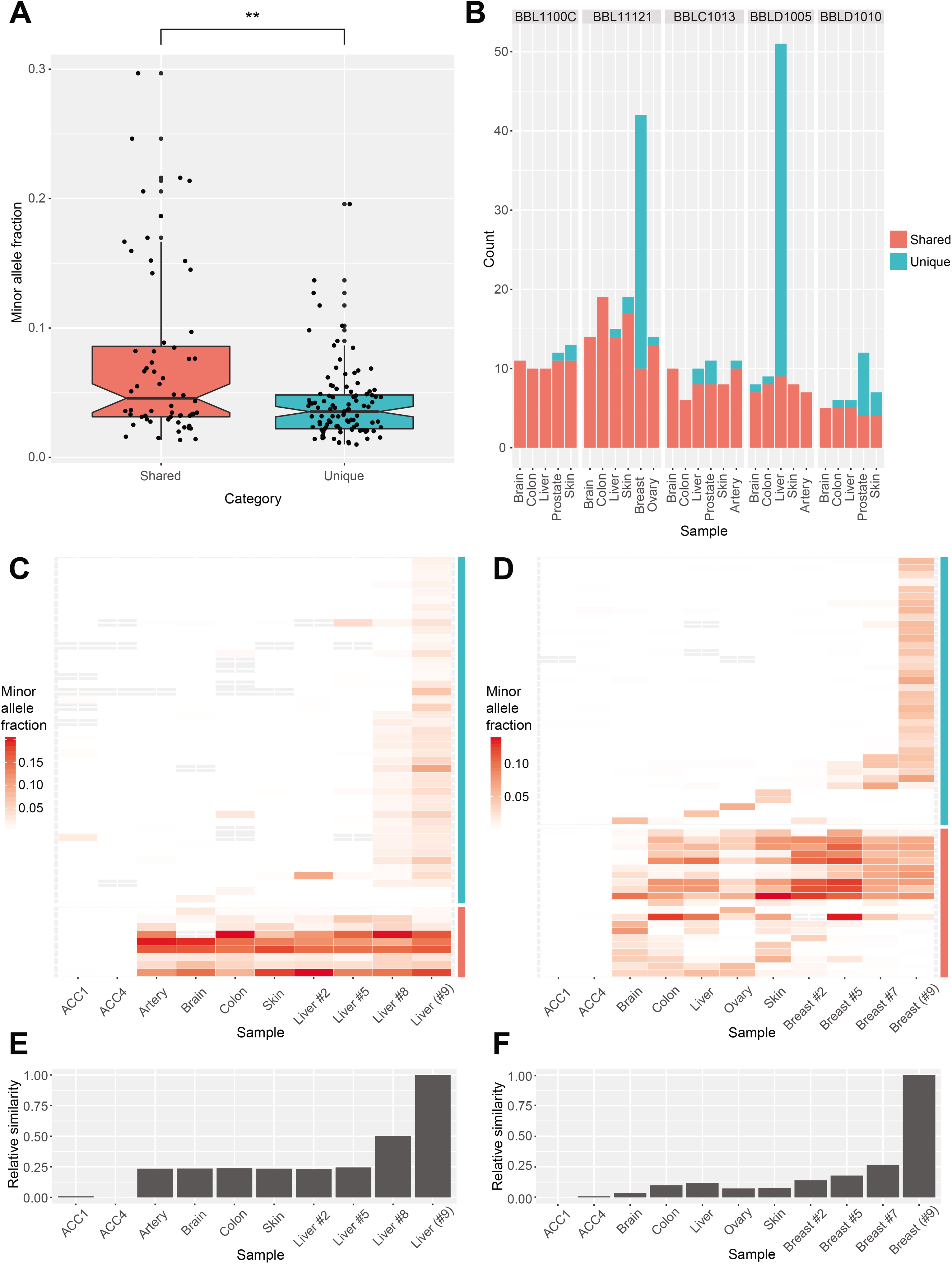
Two types of pSNMs revealed by inter- and intra-organ profiles. (A) Minor allele fractions between different categories of pSNM. The organ-shared pSNMs demonstrated significantly higher allele fractions than the organ-unique pSNMs. (B) Number of pSNMs carried in different organ samples. Red and blue bars denote the organ-shared and organ-unique pSNMs, respectively. An excess of organ-unique pSNMs was observed in the breast sample of BBL11121 and the liver sample of BBLD1005. (C-D) Heatmap of minor allele fractions for pSNMs carried in multiple organ samples of BBLD1005 (C) and BBL11121 (D). Blood samples of two unrelated individuals (ACC1 and ACC4) served as negative controls. The color intensity of each tile represents allele fractions estimated by targeted ultra-depth resequencing. Gray tiles denote sites without sufficient read depth (< 30X). Red and blue bars denote the organ-shared and organ-unique pSNMs, respectively. The majority of organ-unique pSNMs were locally restricted to one or a few physically adjacent organ samples (<1 cm). (E-F) Relative intra-organ similarity of pSNM profiles in BBLD1005 (E) and BBL11121 (F). The originally sequenced samples (liver #9 and breast #9) shared the largest similarity to their physically closest samples.

Close inspection of the pSNMs unique to only one organ revealed a distinctive type of pSNMs. Two organs had a dramatic excess of organ-unique pSNMs compared to other organs (Fig 2B). Specifically, the liver sample of BBLD1005 and the breast sample of BBL11121 carried 42 and 32 organ-unique pSNMs, respectively, compared to an average of 1.1 organ-unique pSNMs for the other organ samples. This suggested that the majority of pSNMs in these two organs might originate organ-specifically after embryogenesis [30].

To further investigate these excessive organ-unique pSNMs in these two organs, we sampled three additional adjacent samples from each organ with varying physical distances to the original samples used for WGS (**S7 Fig**) and applied PASM to profile the allele fractions of validated pSNMs (**S8 and S9 Figs**). While 8 of the 9 (88.9%) embryonic pSNMs could be detected in all three intra-organ samples (Fig 2C and 2D), consistent with our prediction that these mutations occurred early in embryogenesis, the organ-unique pSNMs manifested with a distinct intra-organ pattern. In BBLD1005, 22 out of 42 (52%) liver-unique pSNMs identified in the original sample (liver #9) were also detected in the physically closest sample (liver #8), whereas only one liver-unique pSNM was detected in the two samples further away (liver #2 and #5) (Fig 2C). Given that the physical distance between liver #8 and #9 was about 0.5 cm and the distance between liver #2/#5 and liver #9 was approximately 3.5 and 2 cm, respectively, these results suggested that the majority of liver-unique pSNMs were locally restricted to a small volume of liver cells. A similar observation was made in the breast samples of BBL11121 that breast #7 shared more pSNMs to breast #9 than breast #2 and #5 (Fig 2D). We reconstructed the inter-sample similarity using the minor allele fractions of pSNMs, and indeed the originally-sequenced liver or breast samples shared the largest similarity to their physically nearest samples (Fig 2E and 2F).

Analysis of minor allele fractions of the liver- and breast-unique pSNMs revealed a single narrow peak for each organ sample (**S10 Fig**), with an average of 3.1% and 4.2%, respectively. Considering that such pSNMs were restricted to a small region within the organ, the narrow peaks likely resulted from clonal expansion events during the process of organ self-renewal that generated a sub-population of cells carrying postzygotic mutations large enough to be detected in bulk sequencing [23]. We refer to these pSNMs as “clonal expansion pSNMs” in the following analyses. Our results demonstrated the presence of clonal expansions in various types of non-cancer organs and highlighted clonal expansion as one of the major sources of pSNMs in clinically unremarkable individuals.

### Mutation Spectrum, Replication Timing, and Chromatin Status Varied Between the Two Types of Postzygotic Single-Nucleotide Mosaicisms

If the embryonic pSNMs arose from early mutations during embryogenesis and the clonal expansion pSNMs arose from more recent mutations during organ self-renewal, they may present different mutational characteristics. To explore this possibility, we compared these mutations in terms of mutation spectrum, replication timing, chromatin status, and selection.

We first studied the mutation spectrum of the two types of pSNMs identified. For embryonic pSNMs, C>T mutations were the most predominant type (65.0%), with a significant elevated mutation rate at CpG sites vs non-CpG sites (Proportion Z-test, *P*-value < 2.2×10^-16^, Fig 3A). The enrichment of C>T mutation at CpG sites could be explained by the spontaneous deamination of 5□methylcytosines (5mC) [20], which has also been reported as one of the most common signatures in cancers [38]. The predominant C>T mutation at CpG sites for embryonic pSNMs were consistent with previous studies of early pSNMs in human [14, 39] and mouse [40]. On the contrary, we observed predominant C>A (39.5%) and T>C (42.4%) mutations for the clonal expansion pSNMs identified in BBLD1005’s liver and BBL11121’s breast samples, respectively (Fig. 3B and 3C). Oxidative DNA damage was one of the major cause for C>A mutations [22], and the higher proportion of C>A mutation in the liver sample might reflect the accumulated oxidative stress of hepatocytes.

**Fig 3.**
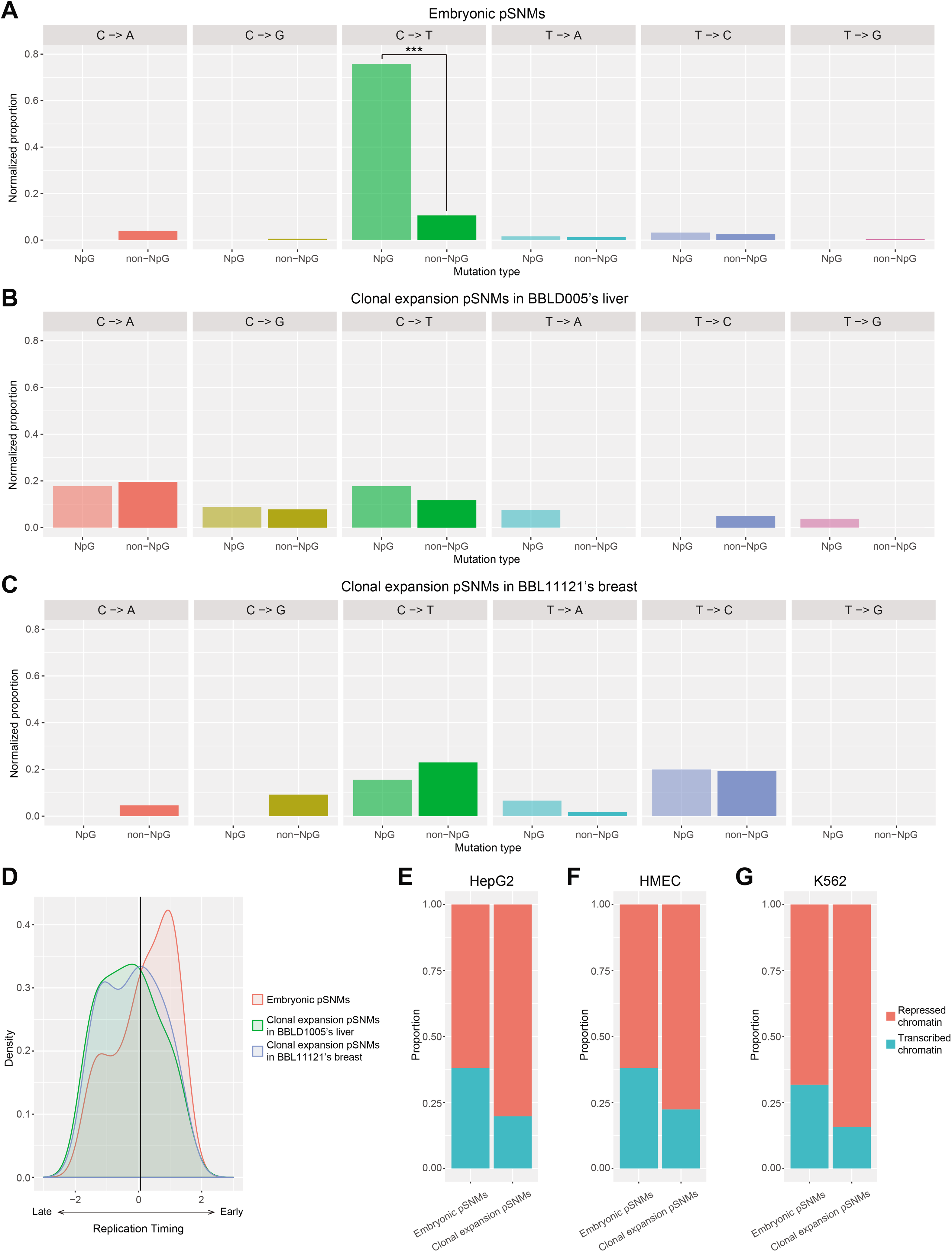
Distinct genomic characteristics between embryonic and clonal expansion pSNMs. (A-C) Mutation spectrums in NpG and non-NpG sites for embryonic pSNMs (A), clonal expansion pSNMs in BBLD1005’s liver (B), and clonal expansion pSNMs in BBL11121’s breast (C). Mutation rate was normalized by the total number of sites in the human genome. For embryonic pSNMs, CpG sites showed significantly higher rate of C>T mutations than non-CpG sites. (D) Varied DNA replication timing of the two pSNMs types. The grey line denotes the genomic average. Embryonic pSNMs were enriched in early-replicating regions, whereas clonal expansion pSNMs were enriched in late-replicating regions. (E-G) Proportion of pSNMs locating in open or closed chromatin regions in the HepG2 (E), HMEC (F), and K562 (G) cell-lines. Significantly higher proportions of embryonic pSNMs were observed in transcribed chromatin regions of all three cell types.

Previous studies had reported elevated rates of germline and cancer-related somatic mutations in late-replicating regions [41, 42]. Using data from the replication timing profile of lymphoblastoid cell-lines [43], we observed significantly different distributions of replication timing between embryonic and clonal expansion pSNMs (Wilcoxon rank-sum test, *P*-value = 9.7×10^-3^; Fig 3D). Clonal expansion pSNMs were significantly enriched in late-replication regions (Permutation test, *P*-value = 0.006), similar to previous reports of germline and cancer-related somatic mutations, while embryonic pSNMs were significantly enriched in genomic regions that replicated earlier (Permutation test, *P*-value = 0.026). Embryonic pSNMs with a wide range of allele fractions contributed to the early-replication enrichment (**S11 Fig**), suggesting that the enrichment was not caused by a small number of outliers. This bimodal distribution was confirmed using the replication timing profiles from five other cell-lines: GM12878, K562, HeLa-S1, HepG2, and HUVEC (Wilcoxon rank-sum test, *P*-value < 0.05). We further confirmed our finding by using the pSNMs identified from the single-clone sequencing of neuronal progenitor cells [14], where the mutations which were shared by other brain regions and non-brain tissues were significantly enriched in early-replicating regions than those specifically present in the clone of neuronal progenitor cells (Wilcoxon rank-sum test, *P*-value = 0.028). The distinct pattern of replication timing between the two types of pSNMs might reflect different mutational effects of replication timing during different stages of human development.

Last but not the least, we investigated whether chromatin status contributed to the mutation rate of pSNMs. For embryonic pSNMs, the genomic distance between a pSNM and its closest DNase sensitive zone in embryonic stem cells was significantly smaller than the expectation under uniform distribution (Permutation test, *P*-value = 013). In contrast, clonal expansion pSNMs did not showed the enrichment of DNase sensitive zone (Permutation test, *P*-value = 0.54). We further found that embryonic pSNMs were significantly enriched in the topologically associating domains (TADs) containing embryonically-transcribed genes (Fisher’s exact test, *P*-value = 0.046), and this pattern was robust with different thresholds for embryonically-transcribed genes (**S12 Fig**). Moreover, we observed a significantly larger proportion of embryonic pSNMs compared to clonal expansion pSNMs within transcribed chromatin regions using epigenetic data from three cell-lines of different origins (Fisher’s exact test, *P*-value < 0.05; Fig 3E-G). Analyses of tissue-shared pSNMs versus clone-specific pSNMs previously identified in neuronal progenitor cells [14] further confirmed our finding (Fisher’s exact test, *P*-value < 0.01; **S13 Fig**). In summary, we reported an elevated rate of postzygotic mutations in open and transcribed chromatin regions during embryogenesis, which might result from the exposure of external or internal mutagens within these regions [44].

## Discussion

Recent researches have significantly expanded what is known about the functional roles of postzygotic mutations, which now include not only cancers and overgrowth disorders [45], but also other complex disorders [11, 46]. With the help of next-sequencing technologies, postzygotic mutations have been identified and validated in healthy individuals [13, 39], confirming the theoretical predictions that postzygotic mutations are prevalent and every person is a mosaic [23]. However, the number of rigorously validated postzygotic mutations in healthy individuals has been small, which has hindered statistical analyses of their genomic patterns. In particular, little is known about the genomic patterns of postzygotic mutations in the normal development process of healthy human organs.

In this study, we discovered two distinct types of pSNMs, one occurring during early embryogenesis and the other likely to occur during later tissue-specific clonal expansion. Surprisingly, these mutations manifested many distinct features in regard to mutation spectrum, replication timing, and chromatin status, implying dynamic mutational effects across different developmental stages. Unsurprising in hind sight, clonal expansion pSNMs shared many mutational features with previously reported cancer mutations [47], as tumorigenesis is a specialized process involving clonal expansion of cancer cells [28]. Previous studies reported high proportion of C>A and T>C mutations as well as enrichment of late-replicating regions for clonal expansion pSNMs that were identified from skin fibroblasts [48–50], which was concordant with our findings of clonal expansion pSNMs in the liver and breast samples. In contrast, our embryonic pSNMs demonstrated a range of unique features, including an elevated C>T mutation rate in CpG sites, an enrichment in early-replicating regions, and a stronger effect of transcribed chromatin status (Fig 3). Similar patterns in mutation spectrum (**S14 Fig**), replication timing (**S15 Fig**), and chromatin status (**S3 Table**) could be observed between the embryonic pSNMs that were globally present in all the sequenced organs of the donor and those only present in some but not all the sequenced organs. To further cross-validate our findings, we further analyzed an independent pSNM list that had been identified from human neuronal progenitor cells [14], and confirmed the varied genomic patterns between pSNMs which originated at different developmental stages (Results). In addition to WGS, the elevated C>T mutation rate in CpG sites was also reported in high-fraction pSNMs identified from whole-exome sequencing data [11, 46].

Two of the reasons why the study of postzygotic mutations in healthy organs lags behind that of tumors include the lack of matched control samples in healthy individuals and the significantly lower abundance of postzygotic mutations. Our results showed that 38% of the validated pSNMs were shared by more than one organ, proving the importance of using a control-free pSNM-caller such as MosaicHunter. Furthermore, the high specificity of MosaicHunter compared to other callers enabled us to generate a candidate list that was specific enough to be validated. Single-cell sequencing has been demonstrated to be an alternative approach to study postzygotic mutations [14, 15]. However, compared to single-cell sequencing, bulk sequencing is able to not only provide the genomic location of the postzygotic mutations but also their allele fractions (Fig 1B), which are informative for assessing the proportion of cells that carry the mutation as well as reconstructing the lineage similarity across multiple samples within an individual (Fig 2E and 2F).

The list of pSNMs that had arisen locally during clonal expansion events in the liver and breast samples identified in our study deserve further discussion here. A cell clone with fitness advantage can predominantly proliferate faster and drive all private mutations that were originally carried by that clone to higher allele fractions, allowing them to be detected by bulk sequencing [26]. Because early embryonic pSNMs might affect only a fraction of cells in a certain organ, clonal expansion events could, in theory, make some pSNMs become undetectable from bulk sequencing if the carrier clone was out-competed. Indeed, we observed the breast #7 and breast #9 samples of BBL11121 had lost nine pSNMs that were detected in other breast samples and other organs of the same individual (Fig 2D). These results demonstrated the dynamics of allele fraction for pSNMs driven by clonal expansion events in healthy individuals. We further screened 1407 cancer-related genes from BBLD1005’s liver samples using panel sequencing, and identified four more pSNMs with allele fraction around 1% (**S4 Table**). However, none of the clonal expansion pSNMs had been previously reported in cancer studies and more functional experiments might be required to examine their relationship with clonal expansion.

The current ~100X WGS bulk sequencing data in our study might not provide enough sensitivity to detect the whole spectrum of pSNMs, especially for those with allele fractions less than 1%. The genomic pattern we reported here were based on the analysis of eight organ types from five individuals. With reduced cost of NGS technology, we can expect a better-characterized spectrum of pSNMs in more and more organ samples and individuals in the future. A combination of deeper bulk sequencing and single-cell sequencing on the same organ sample could provide additional insights for pSNMs with lower allele fractions or even those present in only one or a few cells. This will enable a better characterization of postzygotic mutations in the human population and shed new light on distinguishing clinically-relevant postzygotic mutations from the genomic background.

## Methods

### Sample Collection and Preparation

Twenty-seven postmortem organ samples from five donors were obtained from BioServe Biotechnologies (Beltsville, MD, USA), with an approved protocol from the Institutional Review Board and informed consent obtained from all participants or their legal guardians (Table 1). The clinical histories of all five donors showed no diagnosis of cancer or other known overgrowth disorders. Each organ sample was dissected into nine pieces (roughly 0.5×0.5×0.5 cm each) perpendicular to its long axis and labeled from #1 to #9 (**S7 Fig**). The peripheral blood samples were obtained from ACC1 and ACC4, two clinically unremarkable individuals of Asian descent, with informed consent and approval by the Institutional Review Board at Peking University. Genomic DNA was extracted using an AllPrep DNA/RNA Mini Kit (Qiagen, Hilden, Germany) after homogenization.

### Whole-genome Sequencing

Genomic DNA extracted from one piece (labeled as #9) of each of the 27 organ samples and was used for WGS and subsequent validation. To reduce potential bias introduced by library preparation, three sequencing libraries were constructed independently from each sample using a KAPA LTP Library Preparation Kit for Illumina platforms (Kapa Biosystems, Wilmington, MA, USA). Size selection was performed for each library with a target insert size of 350-450 bp using a Pippin Prep system (Sage Science, Beverly, MA, USA). Libraries were purified using Agencourt AMPure XP beads (1.0× volume; Beckman Coulter, Brea, CA, USA) and underwent subsequent quality control using a 2100 Bioanalyzer (Santa Clara, CA, USA). Each library was sequenced in one lane on an Illumina Hiseq X Ten platform (Illumina, San Diego, CA, USA) using 150-bp paired-end reads.

### Identification of Postzygotic Single-Nucleotide Mosaicisms

Raw sequencing reads were aligned to the GRCh37 human reference genome using the paired-end mode of BWA [51]. The aligned reads were processed using Picard and GATK [52] for the removal of duplicated and ambiguous reads (mismatch >4), indel realignment, and base-quality recalibration. The average depths of processed reads in each organ sample ranged from 83X to 113X (Table 1). CNVs and indels were called using CNVnator [53] and GATK [52], respectively, and all the involved regions as well as annotated repetitive regions were masked, because these regions were more vulnerable to false positives due to mis-alignment or abnormal copy numbers. To maximize the detection sensitivity of pSNMs, the single-sample and paired-sample modes of MosaicHunter [35] were applied to each organ sample with default parameters of genotyper and filters. For the paired-sample mode, the WGS data of the other organs obtained from the same donor served as paired controls. Candidates with at least 5% mutant allele fractions and 3 reads supporting the mutant allele were considered in our subsequent analyses. We randomly chose 16 candidates that are present in the latest versions of dbSNP [54] or the 1000 Genomes Project [55] to validate. As a result, all of them were genotyped as heterozygous rather than pSNMs, suggesting that they were more likely to be caused due to the large variation of allele fraction in WGS. Therefore, we only considered the candidates absent in both databases for thorough validation below.

### Validation of Postzygotic Single-Nucleotide Mosaicisms

All the candidate pSNMs were analyzed using the standard workflow of PASM, which had been previously benchmarked by pyrosequencing and micro-droplet digital PCR [5, 56]. PCR primers for PASM were successfully designed for 241 out of the 251 candidate pSNMs, with their amplicon lengths ranging between 380 and 420 bps; for a few candidates located in highly homologous genomic regions, two rounds of nested PCR were carried out to achieve higher amplification specificity (**S5 Table**). For one-step amplicons, 35 cycles of PCR amplification was carried out using 2X Ex-Taq Premix (Takara Bio, Dalian, China). For nested amplicons, additional 15 cycles of first-round amplification and 25 cycles of second-round amplification was carried out to capture the target region specifically.

Amplicons from different organ samples and individuals were barcoded during library preparation and then pooled and sequenced using either the Ion Torrent PGM or Ion S5 XL (ThermoFisher, Guilford, CT, USA), following the manufacturer’s protocols. Sequencing of the same PASM library on the PGM and S5 XL platforms showed a similar distribution of depth-of-coverage (**S16 Fig**). In each sample, we calculated the 95% credible intervals of minor allele fractions for each candidate site with at least 30X depth [5]. Candidates with 95% credible intervals between 0.5% and 40% were considered a mosaic genotype, whereas those sites with a 95% credible intervals’ lower bound below 0.5% or an upper bound above 40% were considered a homozygous or heterozygous genotype, respectively. Five candidate sites without sufficient depth in the negative control samples were removed. As shown in **S17 Fig**, the inter-organ and intra-organ variation of the minor allele fractions for pSNMs could not be explained by the technical variance induced by DNA extraction or PASM.

The differences in pSNM profiles across multiple samples from one individual were assessed using the Euclidean distance (*D*) of the square root of the allele fraction for all the validated sites. The relative similarity (RS) between sample i and j was defined as

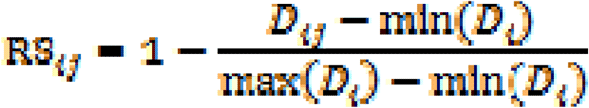

### Copy Number Estimation of Postzygotic Single-Nucleotide Mosaicisms

CNVkit [57] was applied to estimate copy number from the WGS data, and the 100 bp windows centered by each pSNM were considered. The individual-specific copy number was then normalized using the mean across all the five individuals. Three of the 167 PASM-validated pSNMs were found to demonstrate abnormal copy number (< 1.7 or > 2.3 for autosomes and females’ X chromosome and < 0.9 or > 1.1 for males’ X chromosome) in the corresponding carriers (**S1 Table**). The estimated copy numbers were 2.79, 2.80, and 2.59 for these three pSNMs (Q7, P35, and N26), respectively. Their mutant alleles were globally present in all the sequenced organs of the carrier, and the average allele fractions were close to 1/3 (26.2% to 30.9%). Rather than postzygotic mutations, these three sites were very likely to be explained by the germ-line events of copy number gain which made the allele fraction of involved heterozygous mutations deviate from 50%. Therefore, we excluded these three sites from all the following analyses.

### Genomic Annotation of Replication Timing, DNase Sensitivity, and Chromatin Status

The genome-wide annotation of DNA replication timing was extracted from two independent studies [43, 58]. For Hansen *et al.*, we downloaded the wavelet-smoothed signal datasets of five different cell-lines, including GM12878, K562, HeLa-S1, HepG2, and HUVEC [58]. For Koren *et al*, the genome-wide profile was averaged from six lymphoblastoid cell-lines [43]. In both studies, a higher value represented an earlier DNA replication timing. For the permutation test, because our MosaicHunter pipeline only considered candidate pSNMs in non-repetitive regions, we compared the observed median of replication timing for each type of pSNMs against the null distribution estimated by using 1000 times of genome-wide random shuffling among non-repetitive regions.

The DNase-seq data of the embryonic stem cell-line H1-hESC was downloaded from the ENCODE project [59], and the DNaseI sensitive zones were identified using the HotSpot algorithm [60]. For the permutation test, we randomly permutated the genomic positions of pSNMs among non-repetitive regions and assessed the median distance between each permutated pSNM and its closest DNaseI sensitive zone. Each permutation was replicated 1000 times to estimate the distribution under the null hypothesis.

The annotation of chromatin states in HepG2, HMEC, and K562 cell-lines was downloaded from the UCSC Genome Browser [61] to represent cell types derived from the three primary germ layers in early embryo. Chromatin states were inferred from ChIP-seq data of ten epigenetic factors using a Hidden Markov Model [62]. The inferred chromatin states of active promoter, enhancer, and transcription were defined as “transcribed” chromatin status, whereas the other states involving repressed elements and heterochromatin were defined as “repressed” chromatin status.

The annotation of TADs for the H1-hESC cell-line was downloaded from the ENCODE project [59], which was generated based on the Hi-C data of Dixon *et al.* [63]. The expression profile of the same cell-line was also downloaded from the ENCODE project under GEO accession number GSM958733. A gene was defined as embryonically transcribed if its FPKM was larger than 20 in H1-hESC. We also used other FPKM thresholds to confirm the robustness of our finding.

### Cancer Panel Sequencing of BBLD1005’s Liver Samples

To screen for potential driver mutations related to the clonal expansion events in BBLD1005’s liver samples, we captured 1407 cancer-related genes from the liver #2 and #8 samples of BBLD1005 using a Roche SeqCap panel (Pleasanton, CA, USA) designed by Genecast Biotechnology (Beijing, China). The captured libraries were sequenced by Illumina Novaseq6000 (Illumina, San Diego, CA, USA) using 150-bp paired-end reads, with an average depth of ~2000X. Sequencing reads were aligned to the GRCh37 human reference genome by BWA. The pSNMs were identified using MosaicHunter, and the 95% credible intervals of mutant allele fraction were estimated by the Bayesian model implemented in PASM.

## Data Availability

The WGS and targeted re-sequencing data of this study was deposited in the NCBI Sequence Read Archive under accession numbers SRP100797 and SRP136305, respectively.

## Acknowledgments

We are grateful to Drs. Manyuan Long and Chen Xie for their valuable comments and suggestions. Part of the analysis was performed on the High Performance Computing Platform of the Center for Life Science.

**Supporting Information**

**S1 Fig. Distribution of per-site depth-of-coverage for PGM Amplicon Sequencing of Mosaicism (PASM)**. The average depth was greater than 4000X and more than 90% of the candidate sites were sequenced with at least 200X coverage in each sample.

**S2 Fig. Mutant allele fractions and genotypes of candidate sites in multiple organs of BBL1100C**. ACC1 and ACC4 are two unrelated individuals that served as negative controls. The mutant allele fractions were assessed by PGM Amplicon Sequencing of Mosaicism (PASM). Red, green, and blue colors denote heterozygous, mosaic, and reference-homozygous genotypes, respectively.

**S3 Fig. Mutant allele fractions and genotypes of candidate sites in multiple organs of BBL11121**. ACC1 and ACC4 are two unrelated individuals that served as negative controls. The mutant allele fractions were assessed by PGM Amplicon Sequencing of Mosaicism (PASM). Red, green, and blue colors denote heterozygous, mosaic, and reference-homozygous genotypes, respectively.

**S4 Fig. Mutant allele fractions and genotypes of candidate sites in multiple organs of BBLC1013**. ACC1 and ACC4 are two unrelated individuals that served as negative controls. The mutant allele fractions were assessed by PGM Amplicon Sequencing of Mosaicism (PASM). Red, green, and blue colors denote heterozygous, mosaic, and reference-homozygous genotypes, respectively.

**S5 Fig. Mutant allele fractions and genotypes of candidate sites in multiple organs of BBLD1005**. ACC1 and ACC4 are two unrelated individuals that served as negative controls. The mutant allele fractions were assessed by PGM Amplicon Sequencing of Mosaicism (PASM). Red, green, and blue colors denote heterozygous, mosaic, and reference-homozygous genotypes, respectively.

**S6 Fig. Mutant allele fractions and genotypes of candidate sites in multiple organs of BBLD1010.** ACC1 and ACC4 are two unrelated individuals that served as negative controls. The mutant allele fractions were assessed by PGM Amplicon Sequencing of Mosaicism (PASM). Red, green, and blue colors denote heterozygous, mosaic, and reference-homozygous genotypes, respectively.

**S7 Fig. Strategy for intra-organ multi-sampling**. Targeted ultra-deep resequencing was performed on three additional liver samples (liver #2, #5, and #8) with varied physical distances to the original whole-genome sequenced liver sample (liver #9).

**S8 Fig. Mutant allele fractions and genotypes across multiple liver samples of BBLD1005**. The mutant allele fractions were assessed by PGM Amplicon Sequencing of Mosaicism (PASM). Red, green, and blue colors denote heterozygous, mosaic, and reference-homozygous genotypes, respectively.

**S9 Fig. Mutant allele fractions and genotypes across multiple breast samples of BBL11121**. The mutant allele fractions were assessed by PGM Amplicon Sequencing of Mosaicism (PASM). Red, green, and blue colors denote heterozygous, mosaic, and reference-homozygous genotypes, respectively.

**S10 Fig. Distribution of allele fractions for clonal expansion pSNMs in liver and breast samples**. Each sample showed a single peak for the mosaic allele fraction, suggesting that they originated from clonal expansion.

**S11 Fig. Replication timing for pSNMs with varied allele fractions**. The grey line denotes the genomic average. Embryonic pSNMs with a wide range of allele fractions contributed to the enrichment of early-replicating regions.

**S12 Fig. Enriched embryonic pSNMs in the topologically associating domains (TADs) containing embryonically-transcribed genes**. The X axis denotes different FPKM thresholds to define embryonically-transcribed genes, and the Y axis denotes the odds ratio of enrichment between embryonic and non-embryonic pSNMs. The odds ratios were robustly greater than one with varied FPKM thresholds.

**S13 Fig. Chromatin status of tissue-shared and clone-specific pSNMs that were previously identified in Bae *et al.*** The pSNMs shared across multiple brain regions or non-brain tissues showed significantly higher proportion of transcribed chromatin status than those specifically present in the clone of neuronal progenitor cells.

**S14 Fig. Mutation spectrum for the embryonic pSNMs that were globally present in all the sequenced organs of the individual and those only present in some but not all the sequenced organs**. Both types of embryonic pSNMs showed predominant C>T mutations and elevated mutation rate in CpG sites.

**S15 Fig. Replication timing for the embryonic pSNMs that were globally present in all the sequenced organs of the individual and those only present in some but not all the sequenced organs**. Both types of embryonic pSNMs demonstrated the similar enrichment for early-replicating regions.

**S16 Fig. Similar distribution of depth-of-coverage between two platforms used for PGM Amplicon Sequencing of Mosaicism (PASM)**. The red and blue curves denote sequencing runs of the same library performed using an Ion Torrent PGM and an Ion S5 XL, respectively.

**S17 Fig. Varied mosaic allele fractions across different samples of the same organ and different organs obtained from the same individual**. Sample IDs starting with “b”, “c”, “l”, “p”, and “s” denote multiple samples obtained from brain, colon, liver, prostate, and skin of BBL1100C, respectively. Triple technical replicates of PGM Amplicon Sequencing of Mosaicism (PASM) were performed for the V18 pSNM of each sample. The inter-organ and intra-organ variations are dramatically larger than the variation observed between technical replicates.

**S1 Table. Copy number estimation of candidate mosaic sites**.

**S2 Table. List of the 164 validated mosaic sites identified from five individuals**.

**S3 Table. Chromatin state for the embryonic pSNMs that were globally present in all the sequenced organs of the individual and those only present in some but not all the sequenced organs**.

**S4 Table. Mosaic sites identified by cancer panel sequencing of BBLD1005’s liver samples**.

**S5 Table. Primers used for PGM Amplicon Sequencing of Mosaicism (PASM)**.

